# Abnormal mu rhythm state-related cortical and corticospinal responses in chronic stroke

**DOI:** 10.1101/2024.09.12.612604

**Authors:** Miles Wischnewski, Zachary J. Haigh, Taylor A. Berger, Jonna Rotteveel, Tessa van Oijen, Nipun D. Perera, Sina Shirinpour, Ivan Alekseichuk, Rachel L. Hawe, Alexander Opitz

## Abstract

The motor cortex’s activity is state-dependent. Specifically, the sensorimotor mu rhythm phase relates to fluctuating levels of primary motor cortex (M1) excitability, previously demonstrated in young and healthy volunteers. However, it is unknown whether this observation is generalizable to individuals with brain lesions after a stroke. We investigated the phase relationship between the mu rhythm and cortical excitability by combining real-time processing of electroencephalography (EEG) signals and transcranial magnetic stimulation (TMS) of M1. In 11 chronic subcortical stroke survivors and 12 similar-aged healthy volunteers, we applied TMS to M1 at the peak, falling, trough, and rising phase of the sensorimotor mu oscillation. As outcome measures, we investigated the M1-to-muscle excitability by measuring motor-evoked potentials (MEPs) and local cortical activation by measuring TMS-evoked potentials (TEPs). We found that M1-to-muscle excitability in stroke survivors and older volunteers shows a phase-dependency similar to that in young healthy adults. That is, MEPs were increased and decreased at the trough and peak of the mu rhythm, respectively. However, individuals with stronger stroke-related motor symptoms showed a decreased phase preference. Further, phase-dependency was abolished in the local cortical activity, as measured with EEG, in the stroke-affected hemisphere, in contrast to the non-affected hemisphere as well as either hemisphere in healthy volunteers. Altogether, these results shed light on the state-dependency of motor cortex excitability after stroke. Our results indicate that the strength of phase preference of TMS motor responses could indicate the severity of motor impairment. These results could enable the development of improved TMS paradigms for recovery of motor impairment after stroke.

## Introduction

A large number of stroke survivors experience significant motor impairments that persist long into the chronic stage (>6 months after the incident). While continued movement therapy is the gold standard for treatment, many patients experience a plateau of movement ability.^1^ Additional treatment by applying external motor corticospinal activation through transcranial magnetic stimulation (TMS) has been shown to have a superadditive effect on standard therapies.^2–4^ Since TMS is non-invasive and without significant side effects, it is an optimal add-on tool for movement recovery after stroke.^5^

Identifying the optimal stimulation parameters, including location, intensity, and timing, is crucial for TMS to be effective.^5,6^ In particular, the timing of TMS pulses is paramount, as recent work demonstrated that motor corticospinal excitability dynamically changes in tandem with fluctuating brain states as measured with electroencephalography (EEG).^7^ Targeting time-varying brain states requires real-time EEG signal analysis to inform the timing of TMS. In a real-time TMS-EEG setup, EEG brain oscillations are analyzed online to trigger TMS at a particular time of interest.^8–10^ In young, healthy volunteers, it was found that output from the primary motor cortex (M1) to the hand muscles depends on the phase of the ongoing brain oscillation referred to as the sensorimotor mu rhythm (8-13 Hz).^8,11–16^ Independent research has shown that stimulation at the trough-to-rising phase of this brain oscillation results in 10 – 20% larger muscle output compared to stimulation at the peak-to-falling phase.^8,14^

Another way to quantify oscillation phase-dependent effects of TMS, besides M1-to-muscle excitability, is by investigating TMS-induced local activation changes in the EEG.^17–21^ Such TMS-evoked potentials (TEPs), at least the earliest components, have the advantage that they are less confounded by subcortical and spinal activity fluctuations.^22^ While only a few studies have investigated the effects of the TEP phase-dependency, preliminary outcomes suggest that TEP component amplitude may depend on the mu phase.^23^

While such findings highlight the promise of phase-specific stimulation for the clinical use of TMS after stroke,^24^ it is important to note that the above studies were all performed in young, healthy volunteers. It is currently unknown how motor cortex excitability depends on the mu oscillation phase in healthy older adults and stroke survivors. Furthermore, it is unclear how lesion size and motor impairment affect brain rhythm-dependent motor-corticospinal activation. In this study, we characterize the effects of the mu rhythm phase on cortical excitability in chronic stroke patients, as well as in similar-aged healthy volunteers.

## Methods

### Participants

The study included a sample size of N = 23, consisting of N = 11 chronic subcortical stroke patients (mean age ± SD: 57.7 ± 11.4; for details, see supplementary table 1) and N = 12 healthy volunteers (mean age ± SD: 60.6 ± 8.5). Inclusion criteria for all participants were 1) the absence of psychiatric disorders, 2) the absence of ferromagnetic or electric objects in the body, and 3) no pregnancy. Further, for stroke survivors, the following additional inclusion criteria were formulated: 1) A single ischemic or hemorrhagic stroke six or more months before study enrollment; 2) impaired unilateral movement impairment in the upper extremities, 3) no use of botulinum toxin or muscle relaxants in the target extremity in the six months prior to enrollment; 4) normal cognitive capabilities and ability to understand and give informed consent.

The sample size was based on a power analysis (G*power) and results from our previous studies using the same methodology^14^. Notably, while the number of participants is limited, we obtain a large data sample per individual. Typical research studies using TMS collect 10 to 30 samples per condition^25^, while in this study, we obtained 150 data points per condition, resulting in low variance per person. This is reflected by the strong split-half correlation observed in our previous work (r = 0.82)^14^. Considering these data and assuming a medium-to-large effect size (Cohen’s d = 0.65) with a type I error probability of 0.05 and power of 0.8, the suggested sample size was N = 10.

### Motor assessment

An upper extremity Fugl-Meyer assessment (herein referred to as FMA) was performed by trained personnel to assess upper extremity impairment in stroke patients.^26^ During this assessment, the participant performs a set of upper extremity movements to test the ability to move within and outside synergy patterns, explicitly examining the ability to isolate shoulder, elbow, wrist, and hand movements. Each task of the FMA is scored on a scale of 0 to 2 (0 = is unable to perform the movement; 1 = can partially perform the movement; 2 = is able to perform the movement), for a total of 66 points.

### Neurological assessment

Before investigating phase-dependent cortical responses, all participants underwent a magnetic resonance imaging (MRI) scan in a separate session. These scans were performed using a Siemens PRISMA 3T scanner (Siemens Healthineers, Erlangen, Germany) at the University of Minnesota’s Center for Magnetic Resonance Research (Minneapolis, MN, USA). A 32-channel anterior-posterior head coil was used in all scans. T1-weighted MRI (T1w; 0.8×0.8×0.8mm, TE=2.2ms, TR=2400ms) and T2-weighted MRI (T2w; 0.8×0.8×0.8mm, TE=563ms, TR=3200ms) provided lesion identification and quantification in stroke patients and neuronavigation for TMS sessions in both stroke and healthy groups. MRI conversion from DICOM to NIfTI format was performed using Python’s (v3.7.16) dcm2niix command.^27^

### Lesion identification

In order to identify the location and size of lesions, we first employed the LINDA toolkit (v0.5.1) for automated stroke lesion segmentation.^28^ Because LINDA processes left-hemisphere lesions only, the T1- and T2-weighted images for all right-hemisphere lesions were flipped along their left-right orientation using FreeSurfer^29^ prior to LINDA processing. After completing the LINDA pipeline, the segmentation masks were then flipped back to the correct orientation. Following LINDA segmentation, lesion corrections, and refinement were performed by hand in ITK-SNAP.^30^ The corrections were focused on lesion boundary refinement and discontinuity corrections. The final lesion volume was calculated in FreeSurfer.

### Transcranial magnetic stimulation

We used a MagVenture X100 Pro with a Cool B-65 figure-of-eight coil (MagVenture, Atlanta, GA, USA) for biphasic single-pulse TMS over the primary motor cortex (M1). We targeted the hotspot of the left and right first dorsal interossei (FDI) muscle, and the coil was oriented at a ∼45° angle relative to the midline. The BIOPAC ERS100C amplifier (BIOPAC Systems, Inc., Goleta, CA, USA) with a sampling rate of 10 kHz was used to record electromyography and the motor-evoked potentials (MEP) that follow TMS. Self-adhesive, disposable electrodes were placed on the FDI of both hands. To find the motor hotspot, the intensity was set to 60% of the maximum stimulator output, and TMS was applied in a pseudorandom grid around C3 and C4. If no responses were found, the intensity was increased accordingly. The coordinates with the largest MEPs at a given intensity were marked as the motor hotspot in the Brainsight neuronavigation system (Rogue Research Inc., Montreal, Canada). We used this system to track the exact location and coil orientation continuously. We used the adaptive threshold-hunting algorithm from Julkunen^31^ to determine the motor threshold for each hemisphere. The test intensity was aimed at 120% of the resting motor threshold. However, in cases where this intensity was above the maximum stimulator output or uncomfortable for the participant, the intensity was adjusted to a lower value (but always ≥100% of the resting motor threshold).

### EEG with real-time TMS

For EEG recording, we used a 64 active channel cap (10-20 system, actiCAP, Brain Products GmbH, Gilching, Germany) with TMS-compatible electrodes and amplifier (actichamp, Brain Products GmbH, Gilching, Germany). For real-time recording of the EEG data, we used Lab Streaming Layer (LSL). Data was streamed to MATLAB 2022b (MathWorks Inc., Natick, MA, USA) to feed it into the scripts that run our ETP algorithm.^9,14^ Data was recorded with a sampling rate of 10 kHz and was downsampled to 1 kHz. Impedances of EEG channels were kept below 20 kΩ. C3 in the left hemisphere and C4 in the right hemisphere were the electrodes of interest as they are located over the motor hand area. During online phase extraction of the sensorimotor mu rhythm (8-13 Hz), C3 and C4 were referenced to the surrounding 8 electrodes conform a Laplacian method (Left: Fc1, Fc3, Fc5, C1, C5, Cp1, Cp3, Cp5; Right: Fc2, Fc4, Fc6, C2, C6, Cp2, Cp4, Cp6).

The accuracy and validity of our real-time EEG-TMS algorithm have been established previously.^9,14^ This algorithm, the Estimated Temporal Prediction (ETP), consists of two steps. In the first step, before real-time application, the algorithm is trained on data from a 3-minute resting-state recording. This training provides a first estimate of inter-peak intervals of the mu rhythm. In the next step, during real-time application, the inter-peak interval is compared and adjusted, if necessary, to the training data. Based on this, we predict the next peak, fall, trough, or rise phase of the mu cycle is predicted. A trigger was then sent to the TMS device, with a small correction for the processing time of the algorithm (∼8 ms).

The online pre-processing steps are the same for both the training and application phases. Electrodes of interest were C3 and C4 when targeting the left and right FDI motor hotspots, respectively, based on their close proximity. A Laplacian spatial filter was applied to these target electrodes, meaning that the averaged signal of the eight surrounding electrodes was taken as a reference. The data was also temporally filtered, in the mu (8–13 Hz) or beta (14–30 Hz) range, by applying a zero-phase Finite Impulse Response filter.^32^ Data is tracked in real-time using a sliding window (500 ms), and a bandpass brick-wall filter is used to remove signal edges. The common length of the cycle (peak-to-peak interval) is estimated and subsequently validated by simulating the accuracy of phase targeting using the data from the training phase.

During real-time TMS-EEGc we delivered 1200 total pulses, 600 per hemisphere. A 5-minute break was given after every 150 pulses. Throughout the experiment, phases were targeted in a pseudorandom fashion. The inter-pulse interval was jittered between 2 and 3 seconds, after which the next phase was targeted. Participants were shown a nature documentary to maintain consistent alertness throughout the experiment. This documentary did not contain human movements to avoid any mirror movement effects. Experimenters and participants were blinded to the phase condition at any time.

### M1-to-muscle excitability analysis

M1-to-muscle excitability was measured as the electromyographic deflection at the FDI muscle that results from a TMS pulse, known as the MEP. Specifically, the peak-to-peak amplitude in the EMG using a window of 20-60 ms after the TMS pulse was calculated for each trial (Figure 1). Note that at the affected hemisphere of stroke survivors, MEPs of two participants and TEPs of one participant could not be obtained. Trials with excessive background noise in the EMG were flagged if the average of the absolute EMG activity in a 100-ms window before the TMS pulse was 1) above 0.02 mV or 2) larger than the average EMG activity ± 2.5 times the standard deviation compared to resting window at -500 to -400 ms before the TMS pulse.^14,33^ All MEPs were visually inspected to determine if flagged trials needed to be deleted. Note that no automatic deletion was used as EMG background activity can be elevated in stroke survivors due to muscle spasms or increased muscle tone. For healthy volunteers and stroke survivors, 2.4% and 2.9% of trials were removed, respectively. For analysis, a participant’s MEPs were normalized to the overall average of that participant. MEP analysis was performed using MATLAB 2023a.

**Figure 1.**
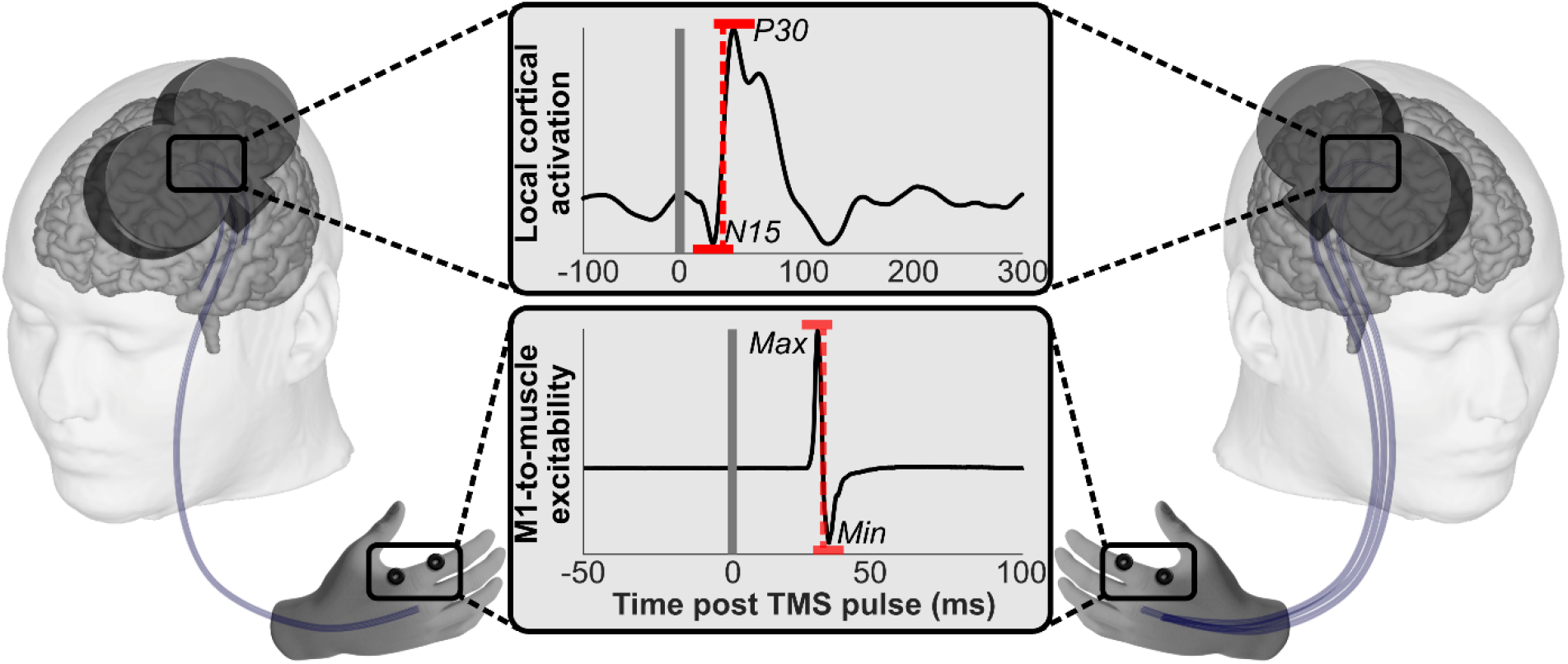
Measurements of M1-to-cortex excitability and local cortical activation. Two outcome measures were investigated. First, M1-to-muscle excitability was measured by investigating the motor-evoked potentials (MEP) amplitude that followed the TMS pulse (lower middle panel). MEPs are observed in the hand (FDI muscle) contralateral to the stimulated hemisphere. Stroke affects the corticospinal tracts, as illustrated on the left panel, with fewer intact connections to the hand muscle (blue curved line, stroke affected) than the right panel (unaffected hemisphere). MEP amplitudes are quantified by calculating the min-to-max EMG difference in a window between 20 and 60 ms after the TMS pulse. Second, local cortical activation was measured by investigating the early components of TMS-evoked potentials (TEP). TEPs are calculated at the EEG electrode closest to the motor hotspot (left hemisphere: C3; right hemisphere: C4). We only investigated the earliest TEP components likely to reflect local cortical activation. Given potential signal offsets in signal due to phase targeting, the second observed component (P30) was subtracted from the first observed component (N15).

### Local cortical excitability

The FieldTrip toolbox, implemented in MATLAB 2023a, was used to analyze local cortical excitability. Specifically, we calculated the deflection in the EEG following a TMS pulse, known as TEP. First, EEG data was epoched into windows starting and ending 1 second before and after the TMS pulse (t = 0). Data within each window was detrended and demeaned. The TMS artifact was removed by interpolating the data from -6 to 10 ms using a cubic Hermite interpolation polynomial. Subsequently, several procedures were performed to clean the data. First, heavily contaminated channels and trials were removed using the FieldTrip artifact summary statistic. Second, an independent component analysis was performed to remove residual artifacts, such as eye blinks and movement artifacts. On average, 3.7 and 3.9 components were removed for healthy and stroke volunteers, respectively. A remaining decay artifact was observed in a small proportion of participants, particularly those in which high TMS intensities were used. Therefore, as a third step, an exponential function was fitted to the first 50 ms of the data.^34^ The exponential function was subtracted from the signal if the fit was larger than r^2^ = 0.4. The signal remained unchanged if no decay artifact was present. Finally, trials were visually inspected to ensure the remaining epochs were artifact-free. Altogether, 6.2% and 9.2% of trials were removed for healthy volunteers and stroke survivors, respectively. After data cleaning, the signal was bandpass filtered between 1 and 50 Hz (3rd order Butterworth) and referenced to a common average. Preprocessing was concluded by using a baseline correction with a window of -500 ms to -15 ms. Finally, the data was averaged per phase (peak, fall, trough, rise), participant group (stroke, healthy), and hemisphere (left, right, or affected, unaffected) to obtain TEPs per category.

Consistent with previous literature, our TEPs consisted of several components: N15, P30, N45, P60, N100, and P180.^17^ Later components are thought to reflect reentrant and network activity, while early components (N15-P30) reflect local M1 excitability directly related to the TMS pulse (Figure 1).^18,35^ Therefore, this article focuses on the early components. We extracted N15 and P30 by identifying the first negative and positive peaks after the TMS pulse. Despite baseline correction, the signal at t = 0 was naturally different for different target phases, which can affect later components. Therefore, local cortical excitability was defined as the difference between P30 and N15, controlling for any signal differences at the time of the TMS pulse. TEP analysis was performed using MATLAB 2023a.

### Statistical analysis

Data was analyzed separately for healthy volunteers and stroke survivors using general linear mixed models (GLMM) with local cortical activation (TEP P30-N15 amplitude) and M1-to-muscle excitability (MEP amplitude) as the dependent variable (Figure 1). The four oscillation phases (peak, fall, trough, rise) and hemisphere (left, right, or affected, unaffected) were used as independent variables. These analyses were repeated with FMA scores as a covariate. The best fitting model (with or without covariate) was determined using the Akaike information criterion (AIC) and was subsequently reported. For stroke survivors, an additional GLMM was performed for the affected hemisphere only, with FMA scores and phase categories as factors. Any significant main and interaction effects were followed by post hoc t-tests. A significance level of α = 0.05 was used for all analyses. We further calculated the individual phase preference (the phase in degrees corresponding to the largest cortical/corticospinal response) and phase modulation strength (the strength of the phase dependency). We calculated the polar vector from the four averaged MEP/TEP values from each phase, where the orientation corresponds to the phase preference and the vector length corresponds to the phase modulation strength. These values were subsequently correlated (Spearman rank correlation) with FMA scores.

## Results

### Lesion volume

The LINDA toolkit successfully identified five out of eleven lesions. The six lesions that the toolkit could not identify were all less than 2300mm^3^; however, low performance at small lesion volumes is a known limitation of the toolkit. For these 6 lesions, the lesion was identified visually and segmented manually. Lesion volume varied widely across participants (mean ± std: 13518.4 ± 31527 mm^3^; Range: 556 - 107408.9 mm^3^). Nine of the eleven lesions included at least a partial overlap with the internal capsule (Figure 2). The other two lesions were localized to the brainstem. Lesion identification for each participant is included in the supplementary material (Supplementary Figure 1).

**Figure 2.**
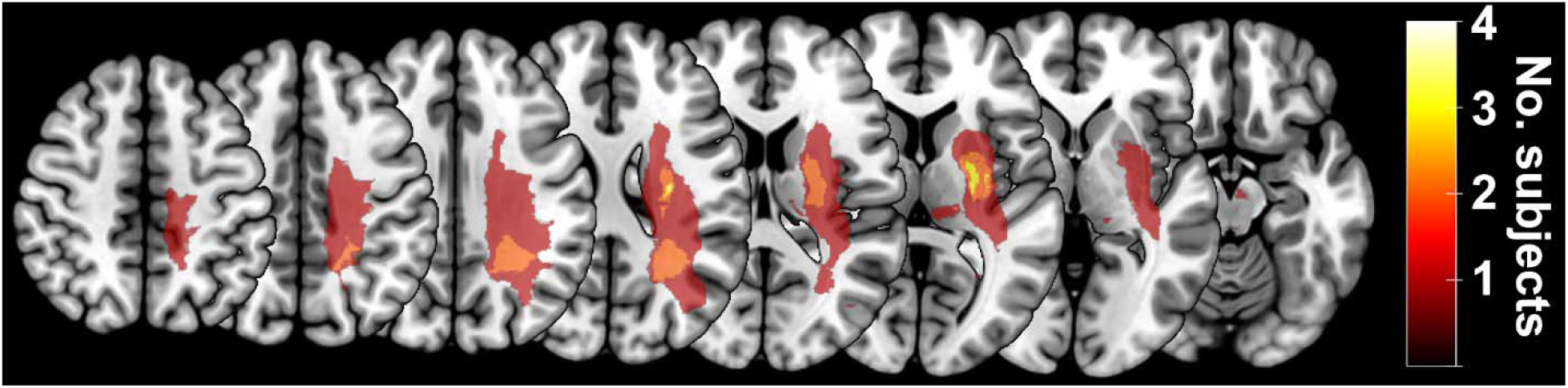
Distribution of stroke lesions. This map represents the overlap in lesion locations across N=11 stroke survivors as displayed on an MNI space T1-weighted image. Lighter colors represent higher overlap, meaning more volunteers have lesions in the same areas. Note that lesions were only shown on one hemisphere by flipping lesions of the right hemisphere to the left.

### M1-to-muscle excitability

Using real-time TMS-EEG, we activated the left and right M1-muscle connection at the peak, falling, trough, and rising phases of the sensorimotor mu rhythm. In individuals with chronic motor disability due to stroke, we found that the mu phase significantly affected M1-muscle excitability. We used a GLMM model with and without FMA scores to statistically test this observation. The latter was a better fit (without FMA: AIC = -243.3, r_adj_^2^ = 0.226, and with FMA: AIC = -236.7, r_adj_^2^ = 0.203) and will be reported here. The results, shown in Figure 3, indicate a significant effect of phase (F(3, 72) = 7.59, p < 0.001). No effect of hemisphere nor a phase*hemisphere interaction was observed (p > 0.47). Consistent for both hemispheres, the highest M1-muscle excitability was observed at the mu trough (affected: 1.044 ± 0.020; unaffected: 1.029 ± 0.011) and the lowest at the mu peak (affected: 0.953 ± 0.017; unaffected: 0.966 ± 0.015). Post hoc t-tests revealed significant differences between peak and trough for both affected (t(8) = 2.57, p = 0.033) and unaffected hemispheres (t(8) = 3.64, p = 0.005). Additionally, comparisons of peak vs. rise (t(8) = 3.63, p = 0.007) and trough vs. fall (t(8) = 3.16, p = 0.013) were significant for the affected hemisphere (Figure 3).

**Figure 3.**
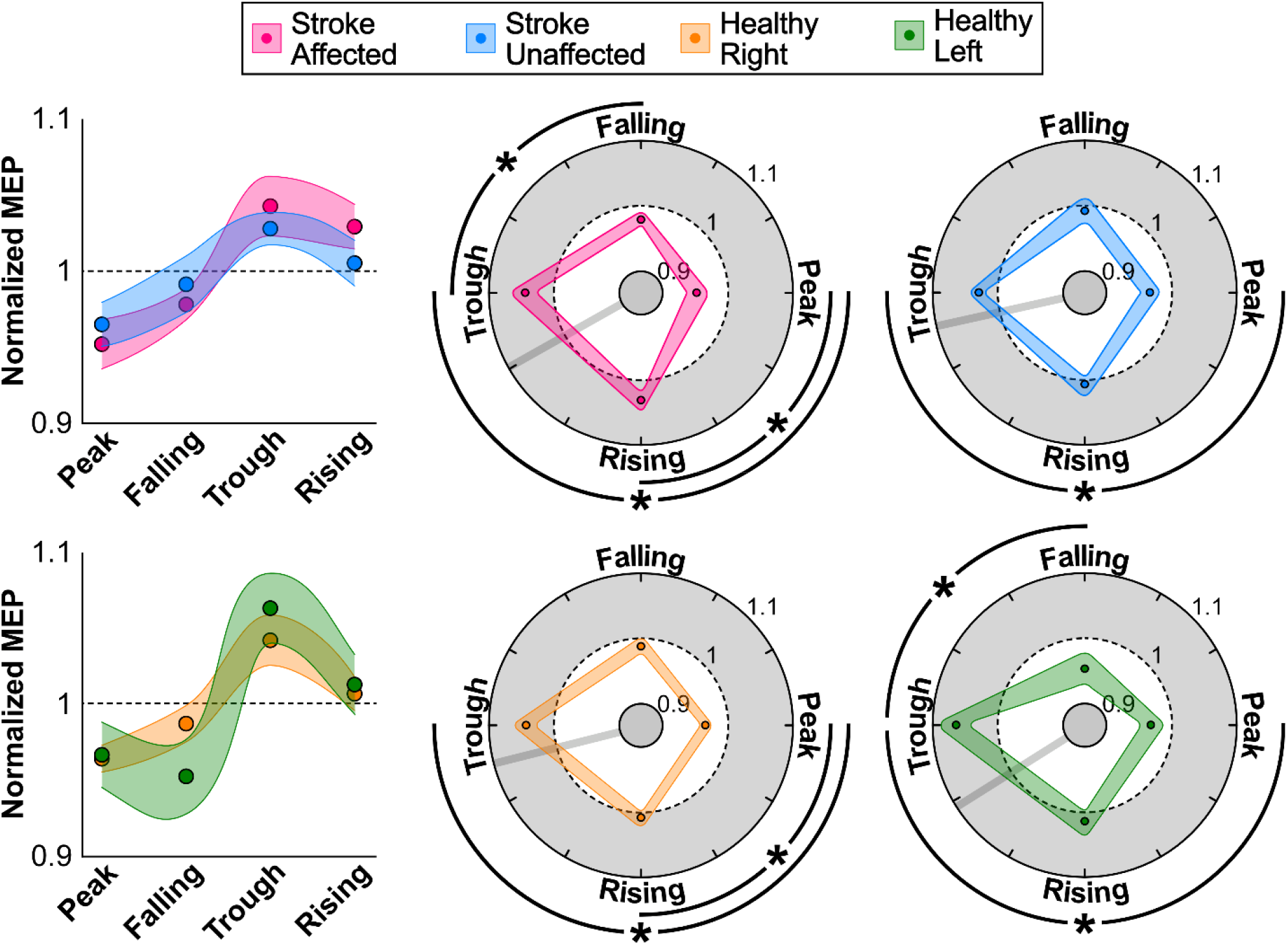
M1-to-muscle excitability per mu oscillation phase. Average ± standard error of normalized MEP amplitudes (M1-to-muscle excitability) for stroke survivors and healthy volunteers, as Cartesian (left) and polar (right) plots. Values larger than 1 reflect increased excitability, while values below 1 reflect reduced excitability. In the polar plots, the preferred phase (direction) is represented by a gray line, and significant pairwise condition differences are represented by an asterisk.

The results were consistent with those observed in the healthy control group, showing a significant main effect of phase (F(3, 92) = 12.99, p < 0.001). Similar to the stroke patients, the largest excitability was observed at the trough of the mu rhythm (left: 1.064 ± 0.023; right: 1.043 ± 0.017), and the smallest excitability was observed at the peak of the mu rhythm (left: 0.967 ± 0.021; right: 0.965 ± 0.018). Post hoc t-tests (Figure 3) revealed a significant difference between the peak and trough for the left (t(10) = 2.89, p = 0.015) and right hemisphere (t(10) = 3.79, p = 0.003).

While FMA was not a significant factor in a combined analysis of affected and unaffected hemispheres in stroke patients, we anticipated that FMA scores are expected to mainly influence the affected hemisphere mainly. Therefore, we also performed a GLMM on data from the affected hemisphere alone. In this case, a model with FMA scores showed the best fit (with FMA: AIC = -110.4, r_adj_^2^ = 0.422; without FMA: AIC = -109.4, r_adj_^2^ = 0.351, respectively). A significant Phase*FMA interaction was found (F(3, 28) = 3.39, p = 0.032), suggesting that the generally observed phase effect depends on FMA scores. To follow up on this finding, we calculated individual phase preference and phase modulation strength per participant (Figure 4), which indicate the angle and the strength of the phase dependency, respectively. We found that phase modulation strength was positively correlated with FMA scores (ρ = 0.750, p = 0.020). This suggests that phase dependency diminishes for patients with a stronger motor disability.

**Figure 4.**
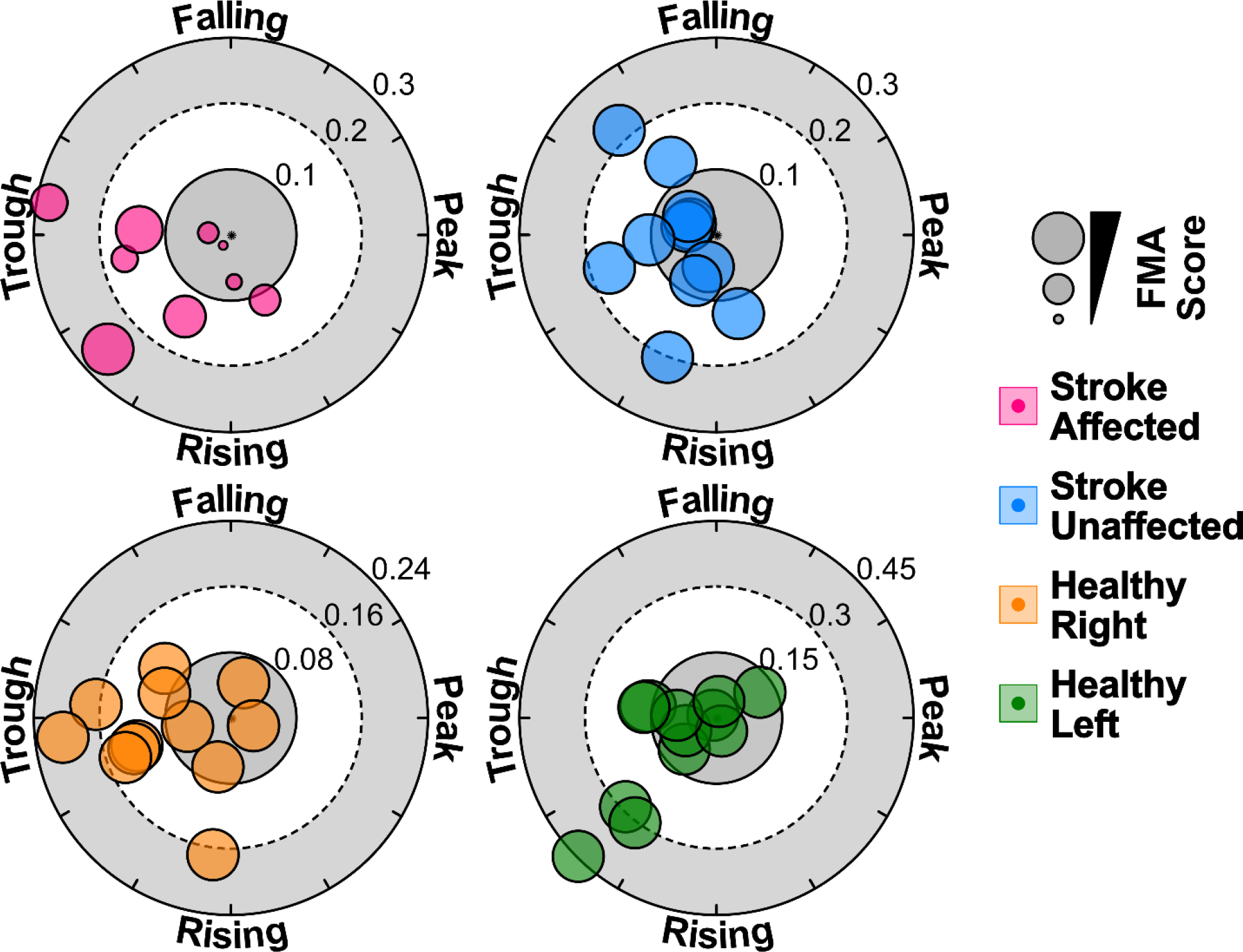
Individual phase preference and modulation strength for M1-to-muscle excitability. For each participant, the polar vector was calculated from the normalized MEP amplitudes of the four phases. The specific angle of the data points represents the phase preference, while the distance from the middle represents the phase modulation strength. Data point size reflects the FMA score, where smaller dots represent stronger movement impairment. Note that the FMA score is only variable for the affected hemisphere, while for the unaffected hemisphere and both hemispheres in healthy volunteers, FMA is maximal.

### Local cortical activation

In addition to investigating the M1-muscle network, we investigated local cortical activation by investigating TEPs at the peak, falling, trough, and rising phases of the sensorimotor mu rhythm. We used a GLMM to investigate the phase-dependency of local cortical activation. We compared models with and without FMA as a factor and found that a model containing FMA was the better fit (without FMA: AIC = -86.58, r_adj_^2^ = 0.042, and with FMA: AIC = -88.75, r_adj_^2^ = 0.113), and hence will be reported here. We found a significant phase*hemisphere interaction (F(3, 71) = 2.81, p = 0.045) with no significant main effects (p > 0.09), as shown in Figure 5. This suggests that the phase effect differed between the affected and unaffected sides. To explore this interaction further, we performed separate GLMMs for each hemisphere. While a significant effect of phase was observed in the unaffected side (F(3, 36) = 7.90, p < 0.001), this was not the case for the affected hemisphere (F(3, 36) = 2.79, p = 0.056). In the unaffected hemisphere, the largest response was observed at the rising phase (1.083 ± 0.036), with the smallest response at the peak (0.919 ± 0.068). A significant difference was observed between the falling and rising phase (t(10) = 2.53, p = 0.030).

**Figure 5.**
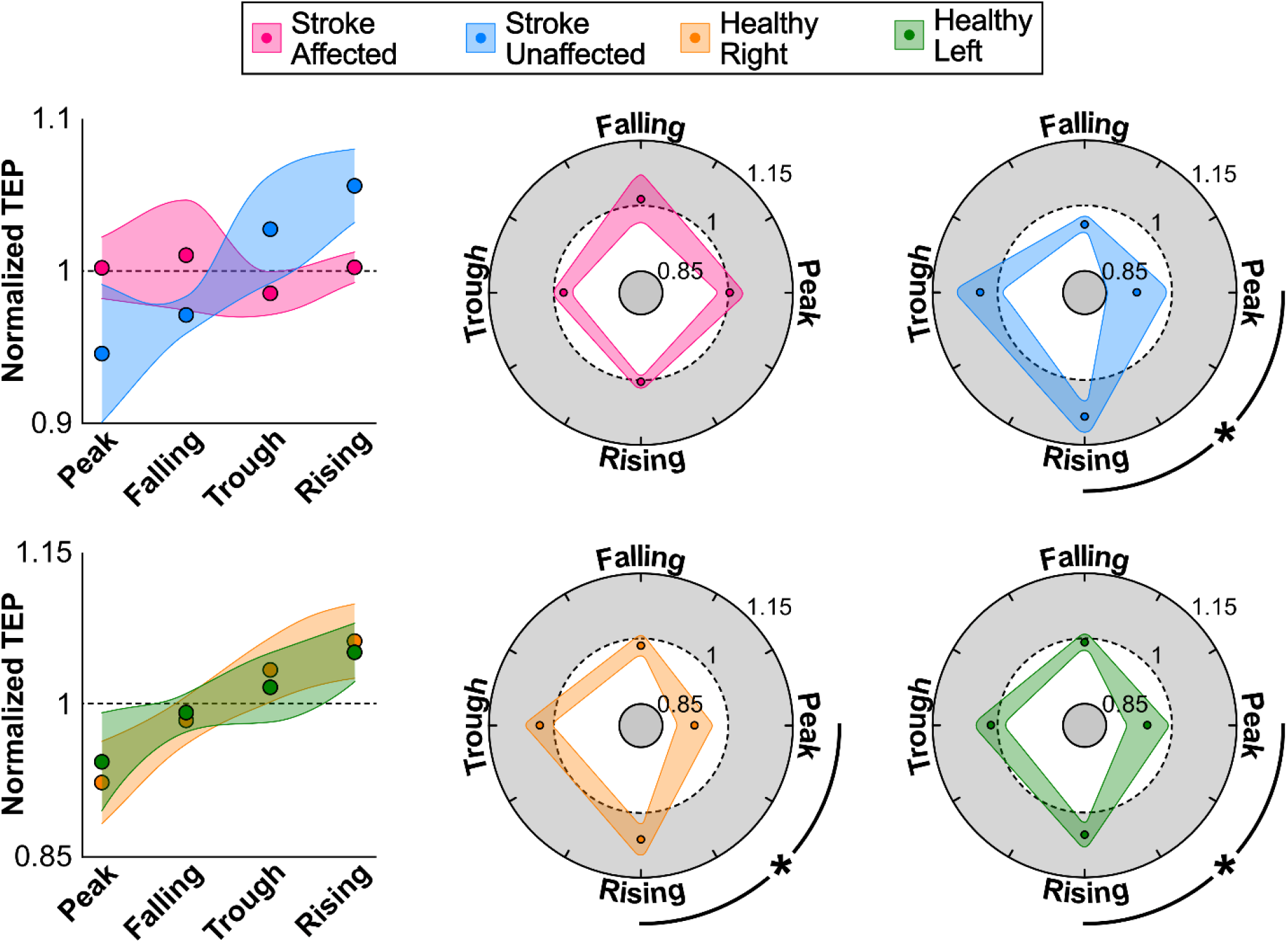
Local cortical activation per mu oscillation phase. Average ± standard error of normalized TEP amplitudes (local cortical activation) for stroke survivors and healthy volunteers, as Cartesian (left) and polar (right) plots. Values larger than 1 reflect increased cortical activation, while values below 1 reflect reduced cortical activation. In the polar plots, the preferred phase (direction) is represented by a gray line, and significant pairwise condition differences are represented by an asterisk.

Furthermore, a significant interaction between phase and FMA was observed (F(3, 32) = 4.26, p = 0.012). However, no significant correlation between FMA and the peak-trough difference (p > 0.1) nor between FMA and phase modulation strength was observed (p > 0.14), as can be observed in Figure 6. This suggests that the severity of motor impairment does not significantly contribute to the effects of phase. In other words, the difference in phase effects between affected and unaffected sides is similar for patients with mild and more severe motor impairments.

**Figure 6.**
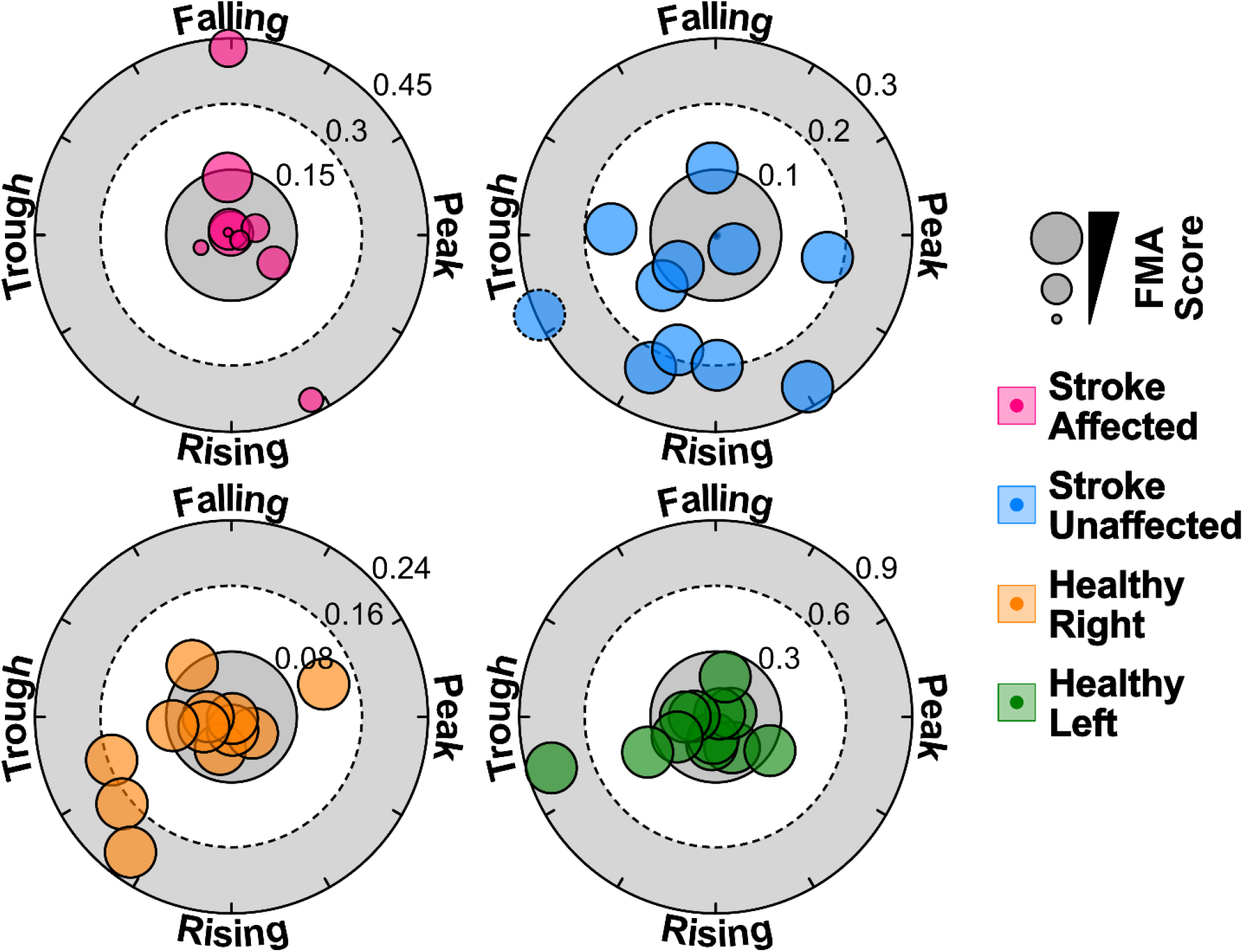
Individual phase preference and modulation strength for local cortical activation. For each participant, the polar vector was calculated from the normalized TEP amplitudes of the four phases. The specific angle of the data points represents the phase preference, while the distance from the middle represents the phase modulation strength. Data point size reflects the FMA score, where smaller dots represent stronger movement impairment. Note that the FMA score is only variable for the affected hemisphere, while for the unaffected hemisphere and both hemispheres in healthy volunteers, FMA is maximal. Further, one outlier was observed in the unaffected hemisphere of stroke patients (blue data point with dotted border) with a phase modulation strength of 1.3.

In contrast to the stroke survivors, healthy volunteers demonstrated no significant interaction between phase and hemisphere (p > 0.56). However, a significant main effect of phase was observed (F(3, 92) = 15.74, p < 0.001), suggesting that local cortical activation depends on phase equally in both hemispheres of healthy volunteers. Post hoc comparisons showed a significant difference between the peak and rising phases (t(11) = 3.02, p = 0.005). In both hemispheres, the largest local cortical activation was observed at the rising phase (left: 1.050 ± 0.028; right: 1.061 ± 0.036), with the smallest responses at the peak (left: 0.943 ± 0.048; right: 0.923 ± 0.040).

## Discussion

This study aimed to characterize oscillation phase-dependent neural responses in stroke survivors and similarly aged healthy human volunteers. Our results point toward several interesting conclusions. First, we found that M1-to-muscle excitability is increased at the trough and decreased at the peak of the mu rhythm in both healthy older adults and the unaffected hemisphere in stroke survivors. This finding replicates previous results in young healthy volunteers.^8,14^ Second, this M1-to-muscle excitability phase-relationship was also observed in the lesioned hemisphere of stroke survivors. However, the correlation with motor assessment scores suggested that this phase modulation strength is smaller in individuals with stronger motor symptoms. Third, local cortical activation, as measured by early TEPs, also showed a phase dependency, with the strongest responses at the rising phase and the smallest responses at the peak of the mu oscillation in the unaffected cortex. Fourth, this phase-dependence of local cortical activation was abolished in the lesioned hemisphere. This absence of phase preference seems consistent regardless of how strong the motor impairment is. Altogether, the results systematically characterize brain excitability depending on the mu oscillatory state in the motor cortex.

The phase of the sensorimotor mu oscillation significantly affected M1-to-muscle excitability in healthy volunteers and stroke survivors. This finding aligns with previous studies in young healthy volunteers that demonstrated the same trend.^8,13,14^ In this study, the difference in normalized MEP amplitude ranged between 6.3% and 9.6%. In a previous study using the same setup, we found a 4.4% phase-induced difference in MEP amplitudes in young, healthy volunteers.^14^ This suggests that the phase-dependent nature of excitability is of the same magnitude or larger in stroke survivors and older volunteers.

The results suggest that the phase-dependency of M1-to-muscle excitability is not significantly changed in a stroke-affected hemisphere. Nevertheless, participants with more severe motor impairment tended to show lower phase modulation strength, meaning brain-state dependency was less pronounced. Lesion size did not significantly correlate with phase modulation strength. The preservation of phase-dependency in the presence of damage provides evidence in support of TMS-based therapies and brain state-based optimization of these therapies. While we do not evaluate therapeutic efficacy in this study, our results may guide future clinical studies.

In contrast, the phase relationship of local cortical activation, as measured by the early TEPs, was abolished at the ipsilesional hemisphere compared to the contralesional hemisphere and the healthy control group. When interpreting the difference in results, it is important to consider that MEPs and early TEPs reflect different aspects of motor activation.^17,21^ Early components of TEPs have been suggested to reflect directly induced activity in the local motor cortical network ^18,35^. Given the extraction of the P30 in this study, this process occurs approximately 30 ms after the TMS pulse.^17^ However, MEPs reflect immediate action potentials of corticospinal M1 neurons about 0-5 ms after the TMS pulse.^36^ Besides the temporal difference, there is also a potential spatial difference. MEPs are thought to originate from the posterior bank of the precentral gyrus.^21^ However, TEPs more likely reflect the activity of the precentral crown, as EEG signals are most sensitive to dipoles oriented perpendicularly to the scalp.^37,38^ Altogether, while TEPs and MEPs reflect early motor activation after TMS, they likely reflect different processes at different times and locations. Therefore, it is not surprising that we found different phase-relation characteristics for both measures.

In this study, only unilateral subcortical stroke lesions were included. This choice was based on the observation that cortical lesions lead to the reduction or absence of the sensorimotor mu rhythm,^39,40^ rendering phase targeting impossible. This aligns with the observation that the mu rhythm has a cortical origin, specifically in the premotor, primary motor, and sensorimotor areas.^41,42^ Therefore, patients with subcortical lesions display a mostly intact mu rhythm.^39,43^ While mu phase-targeting is difficult in patients with cortical lesions, future studies may investigate other approaches, for instance, by extracting the signal from different areas or focusing on alternative brain rhythms.

Furthermore, the present study included stroke survivors in the chronic stage (>6 months post-stroke). In the acute and subacute stages of stroke, several functional and structural changes occur due to neural plasticity,^44^ which may dynamically affect cortical excitability and its relationship with the oscillation phase. In contrast, at the chronic stage, motor improvement tends to plateau,^1,45^ warranting novel treatments to break such a plateau and continue recovery. Repetitive TMS has been suggested as an intervention and has proven successful in improving motor function.^46,47^ However, considerable variability exists, with some studies showing no beneficial effects.^4,48^ To some extent, this variability can be explained by inconsistent outcomes of repetitive TMS.^49^ Recently, it was shown that real-time phase-targeted repetitive TMS overcomes that limitation and leads to more consistent results in cortical excitability.^8,50^ Furthermore, phase-specific repetitive TMS can lead to stronger entrainment of the targeted frequency compared to non-phase-guided TMS.^51,52^ This effect may be further enhanced by entraining oscillations using electrical stimulation.^53,54^

In conclusion, by establishing the mu-phase relationship in local cortical and corticospinal networks and showing a relationship between phase preference and motor symptom severity, we enable future improved clinical TMS studies for stroke recovery.

## Funding

This work was supported by the University of Minnesota’s MnDRIVE Initiative, Brain and Behavior Research Founding (BBRF) Young Investigator grant (29974) and National Science Foundation (NSF) Career Grant (2143852).

## Notes

### Competing Interest Statement

The authors have declared no competing interest.

