## Supplementary Materials for "Abnormal mu rhythm state-related cortical and corticospinal responses in chronic stroke"

| **Supplementary table 1.** Demographics of stroke survivor volunteers. | | | | | | | |
| --- | --- | --- | --- | --- | --- | --- | --- |
| IDX | Type | Age | Sex | PSD (mo) | Dominant hand | Affected hand | FMA |
| 1 | S | 56 | M | 120 | R | R | 53 |
| 2 | S | 47 | M | 141 | L | L | 61 |
| 3 | S | 59 | F | 156 | R | L | 63 |
| 4 | S | 56 | M | 71 | R | L | 18 |
| 5 | S | 71 | F | 38 | R | L | 58 |
| 6 | S | 33 | F | 62 | R | L | 63 |
| 7 | S | 67 | M | 23 | R | R | 20 |
| 8 | S | 50 | F | 17 | R | L | 21 |
| 9 | S | 69 | F | 103 | R | L | 60 |
| 10 | S | 69 | F | 96 | R | R | 57 |
| 11 | S | 58 | M | 25 | R | R | 8 |
| Abbreviations: FMA = Fugl-Meyer Assessment, F = Female, L = Left, M = Male, R = Right, S = Subcortical | | | | | | | |

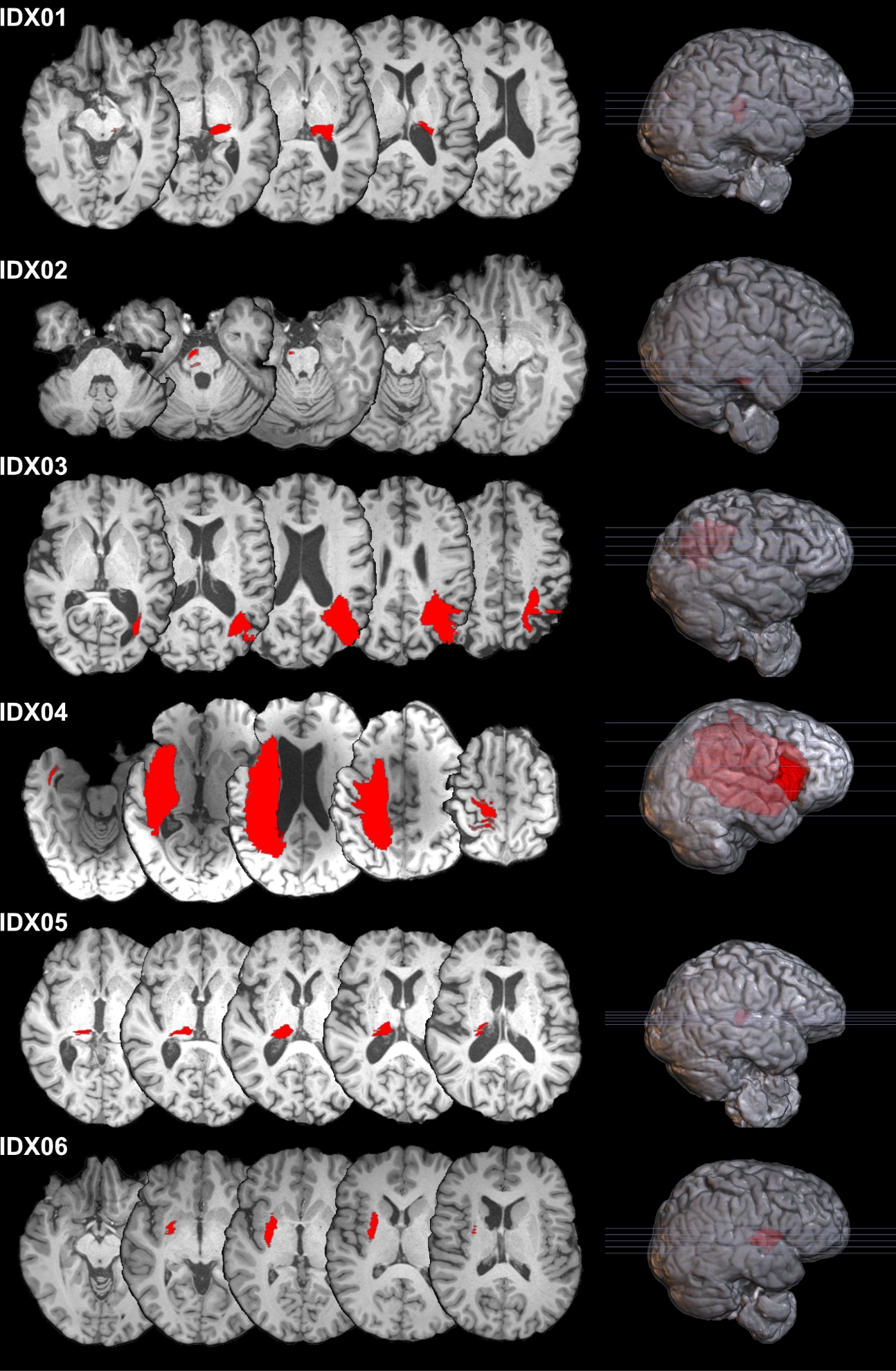

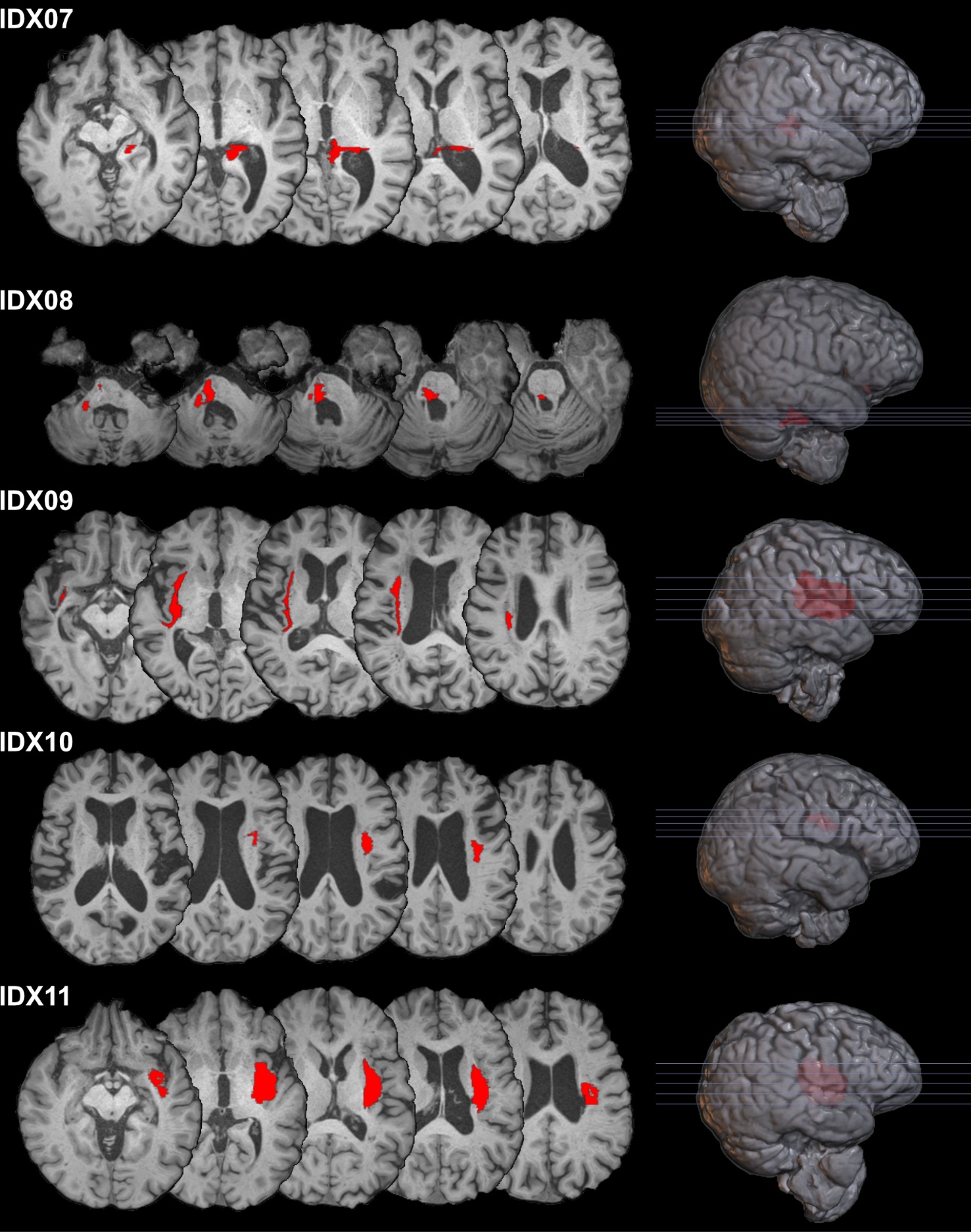

**Supplementary figure 1.** Radiological view of brain and lesion location for all stroke survivor volunteers. Mosaics are individualized to provide five axial slices that best represent each lesion. Lesion location is marked in red.
